# Chemical integration of proteins in signaling and development

**DOI:** 10.1101/015826

**Authors:** Jeffrey M. Dick

**Affiliations:** Language Institute, Chiang Mai University, Chiang Mai, Thailand

**Keywords:** local equilibrium, standard-state properties, chemical affinity, constraints

## Abstract

In developing embryos of eukaryotes, varying concentrations and/or duration of exposure to morphogens induce and repress transcription factors (TFs) in a regular order, thereby acting as signals for cell differentiation. I present a chemical thermodynamic model for the formation and degradation of the morphogen Sonic hedgehog (Shh) and the major TFs involved in dorsal-ventral patterning of the vertebrate neural tube. This two-stage model first considers introduction of Shh into a chemically defined (oxidizing) environment. As the system is driven away from metastable equilibrium, the TFs progressively become more stable than Shh, as indicated by comparison of the calculated overall energies of formation (chemical affinity). In the second stage, a gradual return to metastable equilibrium with Shh drives transitions in the relative stabilities of the TFs that follow the experimental patterns of TF expression. At the intracellular level, the major proteins in the signal transduction network show a metastability progression among reactions of Shh and its cell-surface receptor and the intracellular Gli proteins that act as activators and repressors of transcription. If the endocytosis and degradation of receptor-ligand complexes are coupled to localized protein translation, the consequent thermodynamic constraints may represent a high-level source of specificity that coexists with molecular mechanisms of signal transduction.

## INTRODUCTION

The formation and operation of living systems are made possible only by coupling with a source of energy. In isolation, according to the second law of thermodynamics, a system necessarily departs from an organized, living state. It was recognized by at least the 1920s that many living beings are sustained by the available or “free” energy released by oxidation reactions maintained at non-equilibrium states in the environment (Keller, 2008). The persistence of open, non-equilibrial living systems has been posited as the central observation demanding explanation at the fundamental level in the theory of biology (Scheiner, 2010), and there is continuing interest in biological applications of non-equilibrium thermodynamic models (e.g. Prigogine and Nicolis, 1971; Bizzarri et al., 2013). The outlook on this state of affairs is quite broadly expressed by statements such as “equilibrium is death” (attributed to William Bayliss; see Keller, 2008) and “chemical equilibrium means death” (Jacques Monod, quoted in Judson, 1979).

Does this mean that questions of “where is equilibrium?” and “how far is equilibrium?” are irrelevant to biological studies? This problem is like the difference between explaining how an airplane works using concepts of thrust, lift and drag – questions of dynamics, clearly in the domain of nonequilibrium, and describing how far it is above Earth’s surface, and the bearing to its destination. The latter statements, describing the current state of the plane in reference to its surroundings, are based in equilibrium notions. Equilibrium does not explain dynamic systems, but it does provide a contextual framework for describing certain aspects of organization. One object is “above” another if it has a higher potential energy measured against an externally defined equilibrium position – the ground.

Chemical equilibrium is used as an explanatory concept in some areas of biochemistry, but in many applications it is limited to processes that occur on fast timescales relative to the presumably non-equilibrium changes occurring in the local cellular environment (Garcia et al., 2011). However, there are cases where equilibrium of other types emerges at other levels of organization in living systems. The morphology (form) of organisms is subject to a physical balance of forces, whether they are transmitted directly, or indirectly through heredity (Gould, 1971). For example, physical constraints shape the form of bones; in this and other examples, “equilibrium figures are common in organic nature” as protection from environmental stresses (G. Evelyn Hutchinson, quoted in Gould, 1971). Gould (1980) used “structural integration” to refer to limitations on designs of functional features imposed by physical, architectural constraints.

The concept of allostery in molecular biology holds that the enzymatic properties of a protein, or the strength of binding of a protein factor to DNA, are sensitive to changes in conformation or electronic structure of the protein that can result from “gratuitous” interactions of completely different molecules at different sites on the protein. This disconnect between regulator and substrate has given to molecular biology a guiding principle for signaling and control that is ostensibly free from chemical constraints (Judson, 1979, ch. 10). In contrast, the position taken in this study is that chemical equilibrium, and chemical forces (reaction potential or affinity) have descriptive and explanatory roles in biological systems at higher levels than the mechanism of molecular interactions. This study, which is focused on a higher level of system description, is explicitly *not* a model of molecular interactions. Instead, it will emerge that by utilizing thermodynamic concepts that deal with open systems it is possible to address the effects of subcellular microenvironments on patterns of protein expression important to signaling and cellular differentiation.

There is a growing concern that the conventional mechanistic language of molecular biology is insufficient to account for higher-level developmental and systematic patterns (Kitcher, 1999; Bizzarri et al., 2013). Inherent in the multiple perspectives arising from looking at different levels of organization, the investigation of lower levels is concerned more with mechanisms, while a shift in focus to higher levels reveals context, or constraints, for interactions at lower levels (Pickett et al., 2007). Thermodynamics incorporates a high-level description that guides attention to chemical and environmental constraints. Such a high-level description does depend on abstract simplification, but this is distinct from a mechanistic reduction (Green and Wolkenhauer, 2013). Instead, the thermodynamic model presented here gives predictions about the relative tendencies of protein formation in an local environmental context. This may be found to offer a macro-perspective, complementary to bottom-up models of network interactions based on differential equations, that can help to manage the growing challenges of systems biology (Drack and Wolkenhauer, 2011).

Stability, in terms of equilibrium conditions or steady states^1^, has had a fundamental conceptual role and explanatory relevance to developmental biology. In early embryological studies, cellular determination was seen as a progression through progressively more stable states (Needham, 1942). The different trajectories available to cells along locally stable states (“canalization”) describes the epigenetic landscape metaphor (Waddington, 1956). These models of cellular determination and epigenesis underlie a theoretical definition of morphogenetic fields in the spatial domain of tissues: “Instability (high potentiality) is continually giving place to stability (restricted potentiality)” (Needham, 1942, p. 130). However, a chemical manifestation of these theoretical relationships was elusive; Waddington (1956) felt that biochemistry was not ready “to provide the main framework of ideas for embryology”. More recently, the epigenetic landscape metaphor and morphogenetic field concepts, though still without widely recognized chemical implications, are becoming utilized as useful strategies for system-level descriptions (Balaskas et al., 2012; Bizzarri et al., 2013) and the (re)union of evolutionary and developmental biology (evo-devo) (Gilbert et al., 1996).

In 1948, Sol Spiegelman made an important proposal that unique enzymatic patterns in differentiation arise from a sort of ecological competition for available substrates; as summarized by Gilbert (1996), “The different enzymatic reactions compete for a limited amount of amino acids and energy.” This and related concepts, such as Schoenheimer’s “dynamic equilibrium” of proteins (Judson, 1979), were soon overshadowed by a mechanistic emphasis on persistent, interactive parts that accompanied the rise of molecular biology in the 1950s: “if it were not true that macromolecules, proteins, are stable, molecular biology would not be what it is” (Monod, cited in Judson, 1979). Nevertheless, presently there is little doubt of the continual turnover of proteins, with degradation occurring through both lysosomal and ubiquitin-mediated processes (autophagy and proteolysis) with a wide range of timescales in bacterial and eukaryotic cells (Ciechanover, 2013). Still, there has been very little recent work on the material and energy requirements for macromolecular synthesis and degradation that were recognized by workers in the pre-molecular biology era. Because of recent advances in thermodynamic models and the growing availability of information on protein expression and cellular organization and dynamics, now is a fitting time to re-assess the possible utility of a thermodynamic framework for describing developmental patterns.

In the cellular patterning and differentiation in the development of the neural tube, major regulatory proteins (transcription factors) have spatial expression patterns, connected with a morphogenetic signaling gradient (Jessell, 2000; Ingham and McMahon, 2001; Dessaud et al., 2008; Balaskas et al., 2012). In most descriptions of the intracellular signaling networks, regulation of gene expression (transcription) is explained as a set of interactions between preexisting molecules, most commonly binding interactions (Ingham and McMahon, 2001; Dessaud et al., 2008; Wilson and Chuang, 2010; Hui and Angers, 2011; Cohen et al., 2014). In addition to binding, phosphorylation and dephosphorylation and other modifications (e.g. ubiquitination) are often considered. These different changes lead to the activation or inactivation of the molecules. The signal transduction cascade ultimately leads to the binding of specific proteins (transcription factors) to the DNA, causing changes in gene expression. The analogy of a “relay” was introduced to describe the mediating influence of allosteric proteins (Judson, 1979) and has been also invoked in more recent descriptions of signaling processes (Ingham and McMahon, 2001; Wilson and Chuang, 2010). I refer to models built within this conceptual framework as “relay models”.

While the downstream effects of interactions in the relay models of signal transduction are at the level of the gene (transcription), the steps involved in upstream signal transduction are likely to depend on dynamics of protein synthesis and degradation. The model below takes as its primary focus the forces driving formation and degradation of different proteins, and has the potential to reveal controls on signaling networks, including responses from external stimuli, that act at the level of proteins. I label this description based on the overall requirements of protein degradation and translation as “chemical integration”.

It should be noted that the thermodynamic model itself is oblivious to mechanism; the multiple steps of protein synthesis and degradation are simply not present in the model. However, biological knowledge in the form of proteins that are sequentially induced in tissues or that may be closely localized and reacting in cells form the basis for specifying the starting constraints to build the model. These “top-down” constraints are built into the model as assumptions of local equilibrium among a restricted set of candidate proteins (metastable equilibrium). The thermodynamic calculations give as output a theoretical prediction of the relative stabilities of the proteins that has an explicit dependence on subcellular microenvironmental conditions. The rankings of the relative stabilities of the proteins show which proteins are most likely to form.

In the Methods, a brief survey covers applications of chemical thermodynamics in equilibrium and non-equilibrium systems. In the Results, the model is presented as a thermodynamic framework for describing the observed developmental patterns; the model is based on assumptions about the set of proteins that can chemically react, but it otherwise is not dependent on mechanism. In the Discussion, some speculations are offered on the possible relationships between the thermodynamic calculations and both lower-level molecular mechanisms and higher-level cellular organization. In particular, I suggest that by coupling degradation of endocytosed signaling and receptor proteins with localized translation, the thermodynamic constrains provide a source of specificity that operates in combination with the molecular-level signal transduction and genetic interactions.

## METHODS

By saying “reaction” or “transformation” is meant *only* the change in the abundances of proteins and composition of the system, not the detailed reaction pathway through which the transformation occurs. It is important to keep in mind that a major advantage of equilibrium calculations is not just to consider the final equilibrium state, but to be used as a framework for describing *irreversible* reactions; that is, “equilibrium calculations define limiting conditions for real processes” (Helgeson et al., 1969).

First consider the concepts of *thermodynamic components*, giving a consistent framework for describing compositional differences, and *standard states* and *chemical potentials*, giving a theory for calculating energetic differences. Then, look at the *affinity*, or driving force for reactions that are out of equilibrium. Then summarize extra-thermodynamic considerations, including constraints that can be imposed to describe *metastable* equilibrium.

### Thermodynamic components and reactions

The notion of components is an essential part of a chemical thermodynamic model. The *components* comprise the minimum number of chemical formula units that can be linearly combined to constitute any species (be it mineral, organic species, protein, etc.) in the system under study. Components do not exist in reality, but are defined only in models (Anderson, 2005). Any particular choice of components is admissible as long as is it is one of the minimum number of independent compositional variables in the system (e.g. Helgeson, 1970). The implication is that a choice of different sets of components will contain different variables, and a different numerical description of the system, that nevertheless is completely equivalent in terms of the thermodynamic relations. Accordingly, the actual identities of components is governed by convenience, and is independent of actual structure or “internal constitution” (Gibbs, 1875, p. 117).

The considerations for choosing components may include the ease of making diagrams and interpretations, for example by selecting variables permitting intuitive representations of oxidation state and solubility (Helgeson, 1970), and availability of thermodynamic data for actual species having the formulas of the components (Thompson, 1959). Once the components are chosen, the coefficients in a reaction to form any species from the components are uniquely determined. This also means that, in general, these reactions are not representative of the mechanism of synthesis.

Where the major elements in the system are CHNOS, inorganic species such as CO_2_, H_2_O, NH_3_, H_2_S and O_2_ can be used as components. If a charged species, for example H^+^ is included, the set no longer constitutes thermodynamic components (since phases are electrically neutral), and are referred to as *basis species* (Anderson, 2005). Using basis species, it is possible to consider pH or Eh as independent variables. In this study neutral species are used.

Let us consider the overall formation reaction of a protein from components or basis species. This reaction represents the mass-balance requirements for formation of the human Sonic hedgehog protein (Shh) in a system described by the components identified above.

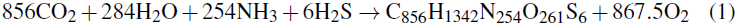

This reaction does not include ionization, phosphorylation, or complexation with metals, which would require other components or basis species. A reaction representing the overall formation of any other protein would involve the same set of components. Again, such a reaction contains no information about the mechanism of synthesis of the biomolecules. Instead, it is a specific statement of the law of conservation of mass. This reaction, or one that could be written for any other molecule or collection of molecules, is the starting point for quantifying the mass and energy requirements for transformations between molecules.

### Affinity and equilibrium

Given the overall formation reaction for any species (Reaction 1), and if the standard Gibbs energies of the species are available, it is possible to calculate the overall “reaction potential” (Helgeson, 1979) or driving force for the reaction, a quantity known as the *chemical affinity* (De Donder and Van Rysselberghe, 1936).

The affinity of a reaction (*A_r_*) is defined as the differential of Gibbs energy of the system (Δ*G*) with respect to reaction progress or extent (*ξ_r_*), at constant temperature (*T*), pressure (*P*) and extent of other reactions (*ξ_k_*) i.e. *A_r_* ≡ − (*∂*Δ*G*/*∂ξ_r_*)_*P*,*T*,*ξ_k_*_ (Helgeson, 1979). The affinity is calculated using (e.g. Helgeson, 1979)

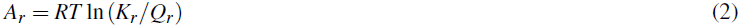

where *R* is the gas constant and *K_r_* and *Q_r_* are the equilibrium constant and the activity quotient of the reaction. *Q_r_* is calculated using

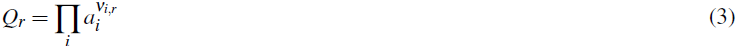

where *a_i_* is the chemical activity of the *i*th species in the system, and *V_i,r_* is the reaction coefficient of the *i*th species in the reaction (negative for reactants, positive for products). The chemical activity is defined according to

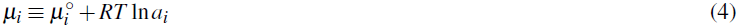

where *μ_i_* is the chemical potential and 
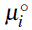
 is the *standard* chemical potential (or standard Gibbs energy, Δ*G*°) of the *i*th species. Eqs. (2)–(4) can be combined to give the familiar expression relating the standard Gibbs energy change of the reaction to the equilibrium constant, 
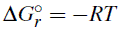
 ln *K_r_*. In Eq. (4), activity (*a_i_*) is replaced by fugacity (*f_i_*) for gaseous species, including O_2_ used in the examples below. In an ideal system, activity can be equated with concentration, so activities (and fugacity for O_2_) are the independent variables used in the calculations here.

The affinity calculated using Eq. (2) is numerically the opposite of the overall Gibbs energy of reaction (Δ*G_r_*; not the standard Gibbs energy, 
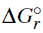
). From Eq. (2) it follows that the affinity is positive if *Q* < *K*, negative if *Q* > *K*, and zero if *Q* = *K*. While *A_r_* is the affinity for a single formation reaction, the calculation of affinities also gives the condition for equilibrium in a system^2^ where the species chemically interconvert (transform). If *A_r_* = 0 for all of the reactions in a system, the system is in *stable* equilibrium; if *A_r_* is equal, but non-zero for all of the reactions, the system is in *metastable* equilibrium (De Donder and Van Rysselberghe, 1936).

The affinity depends on the environment, both through the effects of temperature and pressure on the standard Gibbs energies (affecting *K*), and through the activities of the basis species in the reaction (affecting *Q*). Changes in the latter may occur via reactions internal to the system or may be buffered by the environment. A thermodynamic potential function that is minimized in a system open to one or more components is derived by a Legendre transform of the differential for Gibbs energy, which is used to change the natural variable from moles of components to chemical potentials of components (Alberty, 2003; Anderson, 2005). This is the basis for the theory of “perfectly mobile components” (Thompson, 1959; Korzhinskii, 1965), giving a method for describing the dependence of equilibrium states of *open systems* on chemical changes in the environment.

It is instructive to compute affinities for a number of different species (e.g. proteins) given a set of common environmental conditions and protein activities. Comparing the affinities of the different proteins, the most stable one has the reaction with the highest affinity. Because affinity scales with the size of the molecules, in performing such a stability comparison it is usually desirable to divide all reactions and affinities by the number of amino acid residues (Dick, 2008).

By varying one or more of *T*, *P* or the chemical activities of the basis species, it is possible to investigate the dependence of the relative stabilities of the proteins on these parameters. These relations are conveniently shown on a *predominance diagram*, which shows the most stable species (the one that would be predominant at equilibrium, i.e. have the highest mole fraction) in the system as a function of two variables (on the *x* and y axes). If both of these variables are activities (or fugacities) of the basis species (an *activity diagram*), then the relative positions of the predominance fields and the slopes of the boundaries between them are set by the compositions of the proteins (specifically, the slopes are given by the negative reciprocals of the ratios of the reaction coefficients; Helgeson, 1967). The absolute positions of the boundary lines, however, are set by the standard Gibbs energies of the reactions and the activities of the basis species that are not variable on the diagram.

In the calculations described below, two variables are considered: fugacity of O_2_ and activity of H_2_O. These are chemical potentials in the calculation, and are not constrained in any way other than to explore the effects of their changes on the relative stabilities of the proteins. The numerical values of log *f*_O_2__ and log*a*_H_2_O_ should not be taken to represent actual values (partial pressure or humidity), but rather are *indicators* of the state of the system (Anderson, 2005), that reflect the composition of the system, or the buffering of the chemical potentials by other assemblages of species. The challenge of attaching meaning to very low, unmeasurable values of oxygen fugacity has long been acknowledged by geochemists. A striking demonstration is the calculation showing that an oxygen fugacity equal to 10^−65^ corresponds to one molecule of O_2_ in a volume the size of the solar system (Anderson, 2005). The remark made by Helgeson et al. (1993) is relevant here: “It should perhaps be emphasized in the context of reactions among organic species that *f*_O_2(g)__ does not necessarily refer to free oxygen and that use of log *f*_O_2(g)__ to describe equilibrium relations among these and other species does not require a gas phase to actually be present in the system.” Likewise, *a*_H_2_O_ should be interpreted not as the actual “water activity” but, like *f*_O_2__, as an indicator of the state of the system.

The interpretations from this analysis are often limited to variation in one or two chemical potentials as represented by the diagrams, but in cells there are doubtless many simultaneous changes. The projection on to two dimensions on predominance diagrams, however, helps to formulate hypotheses about the operational range of parameters in the system. Then, it is useful to plot the affinities, or equilibrium distributions, of all of the species along a provisional trend in parameters in order to view the relations of the less stable proteins.

A thermodynamic database and functions to create these and other diagrams for proteins are available in the CHNOSZ software package (Dick, 2008)^3^. The Supporting Information contains the specific code scripts and data files used for the calculations in this paper.

### Metastable and local equilibrium

If some reactions proceed too slowly to reach a stable state a metastable equilibrium among the other species can result. Metastable equilibrium can be considered as a type of partial or constrained equilibrium (Anderson, 2005, 2014) and can be modeled by only allowing certain species to react. Metastable equilibrium also has an energetic definition, as coexistence of species with equal, but non-zero affinities of formation (De Donder and Van Rysselberghe, 1936), so the energy of system is not a global minimum. Metastable equilibrium has been recognized as an important feature of organic molecules in geochemical systems. One example is the apparent metastable equilibrium among organic acids, some hydrocarbons, minerals, and CO_2_ in hydrocarbon reservoirs, explored by Helgeson et al. (1993), who also noted that “mere recognition of a given equilibrium state carries no necessary causal implication with respect to mass transfer processes that may have led to the state”.

The focus above was on global equilibrium states, but the thermodynamic principles can be extended to irreversible systems with the adoption of the *local equilibrium* hypothesis. In a system that is not at equilibrium, the hypothesis maintains that in a small enough volume (“locally”), the entropy and other state variables can be defined (Anderson, 2005). Under this assumption, thermodynamic state variables, which are defined only for chemical equilibrium, can be used to describe and quantify the states and conditions of local systems within a larger non-equilibrium system. For example, local equilibrium in the context of irreversible reactions is the basis for thermodynamic models of metasomatic zonation occurring through transport processes involving introduction or removal of material (Thompson, 1959; Helgeson, 1979), and is also a basic assumption in non-equilibrium theory of entropy production (Prigogine and Nicolis, 1971). Here the conjecture made is that local and metastable equilibrium can be applied to subcellular and multicellular systems containing macromolecules (proteins).

## RESULTS

The development of the neural tube in animal embryos involves complex interactions between morphogens (proteins and other molecules that relay a concentration-dependent signal of positional information), transcription factors (intracellular DNA-binding proteins that regulate gene expression), and other molecules. Dessaud et al. (2008) give an invigorating review of the major features of dorsal-ventral patterning, in which the protein called Sonic hedgehog (Shh) acts as a graded morphogen. To summarize the situation very briefly, Shh is synthesized in cells located at the notochord and floor plate (FP) at the ventral side of the neural tube, and its diffusion and reactions through the tissue results in concentration gradients. Various intracellular transcription factors (TFs) are induced depending on the concentration and timing of exposure of cells to Shh. The specific TFs are responsible for differences in gene expression and the formation of different cell types (neural progenitors) that give rise to e.g. motor and sensory neurons. This complex system has been the subject of studies that consider the physical constraints on diffusion, cross-dependence of activation and inhibition, and computational simulation at the level of gene regulatory networks (see Lander, 2007 for a conceptual review and Cohen et al., 2014 for a recent model of transcriptional regulation). Inherent in the existing models is that morphogens are subject to degradation reactions, as they diffuse or otherwise are transported from the source (Dessaud et al., 2008; Lander, 2007). The major patterns of TFs can be represented by their association with the different progenitor types, as summarized in Table 1.

**Table 1.** A summary of the transcription factors expressed in neural progenitor cells in the ventral half of the neural tube, adapted from Fig. 2B and C of Dessaud et al. (2008). The floor plate (FP) defines the ventral pole of the neural tube; the progenitor domains (p) include the progenitors of motor neuron (pMN) and the most dorsal domains (pD6) that are influenced by Shh signaling. The first 3 columns listing the transcription factors are arranged in order of appearance of Olig2, Nkx2.2 and Foxa2 in the ventral domains. The parentheses represent more detailed trends in the expression patterns (see text).

| Progenitor domain | Transcription factors |  |  |  |  |
| --- | --- | --- | --- | --- | --- |
| pD6 | Irx3 | Pax6 | Pax7 | Dbx2 | Dbx1 |
| p0 | Irx3 | Pax6 |  | Dbx2 | Dbx1 |
| p1 | Irx3 | Pax6 |  | Dbx2 | Nkx6.2 |
| p2 | Irx3 | Pax6 |  | Nkx6.1 |  |
| pMN | Olig2 | (Pax6) |  | Nkx6.1 |  |
| p3 | (Olig2) | Nkx2.2 |  | Nkx6.1 |  |
| FP | (Olig2) | (Nkx2.2) | Foxa2 | Nkx6.1 |  |

### Metastable equilibrium model for Shh as an intercellular morphogen

For the calculations, human protein sequences were taken from the UniProt database (The UniProt Consortium, 2015). The sequence for Shh is that of the 174-residue N-terminal product that is part of the active morphogen. The post-translational modifications by cholesterol addition at the C terminus and palmitoylation at the N-terminus (Dessaud et al., 2008) are not included here. These modifications are likely to have a significant impact on the stability relations, but thermodynamic data for the lipid groups are not yet available and should be considered in future studies. Basis species used are as shown in Reaction 1, with constant activities given by log*a*_CO_2__ = −3, log*a*_NH_3__ = −7, log*a*_H_2_S_ = −7 (using base 10 logarithms); log *f*_O_2__ and log*a*_H_2_O_ are used as exploratory variables. Because no generally applicable mixing model is available, activity coefficients of proteins are taken to be unity; therefore activities are provisionally equated to concentrations, in molal units (mol kg^−1^). The energies of reactions are divided by length (number of amino acid residues) of the proteins; so the lines in the metastable predominance diagram in Fig. 1 represent equal activities of the residue equivalents of the proteins. Likewise, the affinities plotted in Figs. 2–3 are per-residue affinities. Standard Gibbs energies of aqueous unfolded proteins were calculated as described in Dick et al. (2006), with corrected values for the methionine sidechain group, which do significantly alter the results, taken from (LaRowe and Dick, 2012).

**Figure 1.**
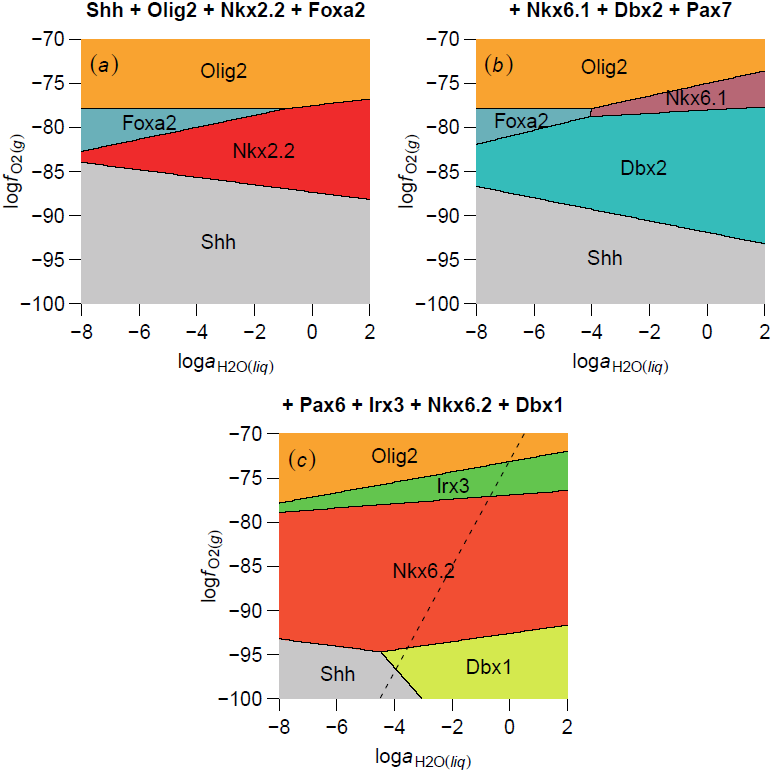
Theoretical metastable equilibrium predominance diagrams for the morphogen Sonic hedgehog (Shh) and selected transcription factors (TFs) involved in dorsal-ventral patterning of the neural tube. The titles show the names of the TFs included in the calculations; in (*b*) and (*c*), the system accumulates the TFs that were present in the preceding calculations. Although Pax7 and Pax6 are included in the calculations for the latter figures, they are calculated to be less stable than the other proteins, and therefore do not have predominance fields. The dashed line in (*c*) shows the trajectory of log*a*_H_2_O_ and log *f*_O_2__ used in Figs. 2 and 3.

**Figure 2.**
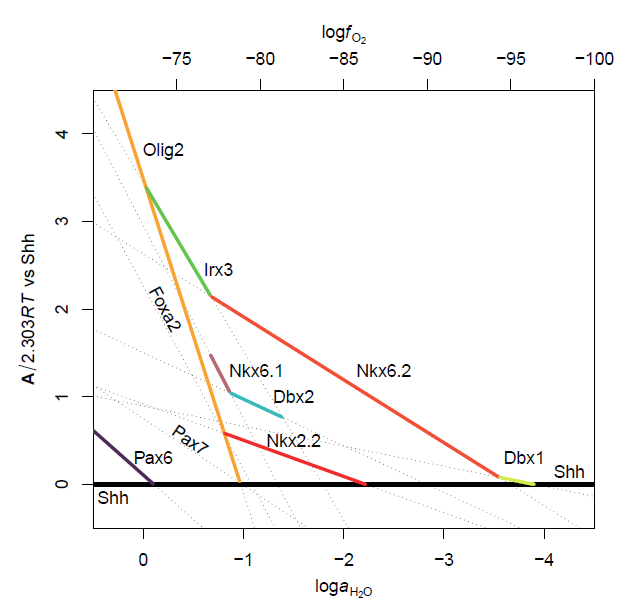
Calculated chemical affinities of the overall formation reactions of Shh and transcription factors (see Table 1). Affinities as a function of changing log *f*_O_2__ and log*a*_H_2_O_ are calculated as per-residue values and are shown as differences from the affinity of Shh. The bold horizontal line represents Shh and the dotted lines with slope < 0 correspond to the various TFs. The colored segments highlight energetic differences that can be related to the developmental patterns (see text and Fig. 3).

**Figure 3.**
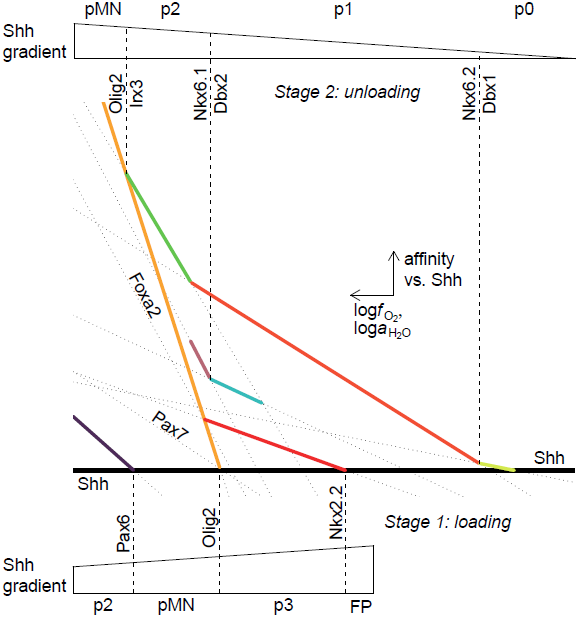
Interpretation of energetic calculations as a framework for describing the linkage between an Shh gradient and appearance of transcription factors during dorsal-ventral patterning of the neural tube. The base diagram is taken from Fig. 2.

The first set of calculations includes the morphogen (Shh) and the most ventral transcription factors. Of these, Olig2 is first formed ventrally, at the FP; with time Nkx2.2, then Foxa2 replaces the ventral expression of Olig2 (Table 1; see also Fig. 2C in Dessaud et al., 2008). Toward more dorsal domains, the expression of Pax6 is increased, giving the progression Nkx2.2-Olig2-Pax6 for the p3, pMN, and p2 progenitors, with a low expression of Pax6 in pMN (Balaskas et al., 2012).

In Fig. 1a is a metastable equilibrium predominance diagram showing the results of the thermodynamic calculations. Shh occupies the more reducing (low-log *f*_O_2__) side of the diagram, while Olig2 is at the opposite end, and the other TFs are in the middle. Accordingly, Olig2 can be most easily formed in oxidizing conditions. Under these conditions, Shh is unstable and has the potential to react. In the second diagram (Fig. 1b), more proteins are added: Nkx6.1, Dbx2 and Pax7. These proteins are active in more dorsal zones of the neural tube (Table 1; Fig. 2 in Dessaud et al., 2008). Pax7 does not appear in the diagram; it is energetically less stable than the others. Nkx6.1 and Dbx2 have replaced (covered up) the stability fields of two of the earlier TFs (Foxa2 and Nkx2.2). The diagram shows that if Nkx6.1 is present, Shh is unstable and has the potential to react. The addition of the last set of TFs (Irx3, Nkx6.2, Dbx1 and Pax6) considered in this model is shown in the third diagram (Fig. 1c). Once again, these new TFs are calculated to be more stable, and their stability fields cover up those of the previous proteins, except for Shh and Olig2.

The diagrams can be used to predict changes in the relative stabilities of TFs resulting from reaction with Shh manifested as changes in the log *f*_O_2__ and log*a*_H_2_O_ values of the microenvironment. For example, in Fig. 1a, reaction of Shh (that is, its irreversible degradation), would bring the system to a more reduced state, leading to a stabilization of Nkx2.2 over Olig2. In Fig. 1b, reaction of Shh favors formation of Dbx2 over Nkx6.1. In Fig. 1c, reaction of Shh destabilizes Irx3, promoting formation first of Nkx6.2, then Dbx1. The directions of the theoretically predicted reactions noted here for Fig. 1b and c are in the *same* order as the observed ventral-dorsal progression (Table 1), but in *reversed* order for Fig. 1a. The significance of these trends is explored in more detail below.

### Intercellular model: energetic interpretation

The diagrams in Fig. 1 show only the metastably predominant protein, that is, the one with the highest affinity per residue under specified conditions. A provisional linear relationship between log *f*_O_2__ and log*a*_H_2_O_ is shown by the dashed line in Fig. 1. By postulating an environmental trajectory that can be projected onto one dimension it becomes possible to graphically compare the affinities of all the molecules rather than show an identification of only the most stable one.

The affinities of formation of proteins are strongly inversely related to oxidation state of the system (e.g. Fig. 6b of Dick, 2014). To visualize the differences between proteins, the affinities in Fig. 2 are plotted as differences from the values calculated for Shh. Therefore, Shh appears as a horizontal line at zero; lines above and below this line correspond to proteins that are more or less stable, respectively, relative to Shh, in the given conditions. Notably, the lines for the TFs all have negative slopes, so that Shh becomes the most stable protein in sufficiently reducing conditions. As log *f*_O_2__ and log*a*_H_2_O_ are increased, the TFs become more stable than Shh in a definite order. Olig2, with the most negative slope, becomes the most stable protein in very oxidizing conditions.

The same affinities are shown in Fig. 3, but with legends added to suggest a thermodynamic description of the distribution of Shh and transcription factors during dorsal-ventral patterning of the neural tube. Among the series of TFs predicted to become relatively stable can be found the sequence Nkx2.2, Olig2, and Pax6 (colored lines intersecting the Shh line at the bottom of Fig. 3). These TFs are required for differentiation of the p3, pMN and p2 domains, in this order (Balaskas et al., 2012).

Suppose that Shh is introduced from the notochord into an environment that is initially at high log *f*_O_2__ and log*a*_H_2_O_, far from the conditions where Shh is relatively stable. A thermodynamic interpretation based on the calculations shown above is that the reaction of Shh (i.e. its degradation and the subsequent alteration of the chemical composition of the local microenvironment) drives down the values of log *f*_O_2__ and log*a*_H_2_O_; more so near the source, where more Shh is available. The lowest values of log *f*_O_2__ and log*a*_H_2_O_ are permissive for formation of Nkx2.2 but not Olig2 or Pax6. Increasing log *f*_O_2__ and log*a*_H_2_O_, that is, a decreasing influence of Shh, is favorable for formation of Olig2, then Pax6 in a stepwise fashion.

The thermodynamic story is, however, more complicated than this. At the conditions where Nkx2.2, Olig2 and Pax6 first become stable relative to Shh, other TFs are even more stable (higher lines in Fig. 3). Thus, the formation of Nkx2.2, Olig2 and Pax6 represent irreversible reactions moving *toward* an equilibrium state, but not arrival at the metastable equilibrium state, which would comprise an assemblage of proteins, with the most abundant being the one with the lowest energy (highest affinity). The appearance of Nkx2.2, Olig2 and Pax6 in the thermodynamic framework can therefore be conceived as a *loading* stage of Shh-driven differentiation, as portrayed in the bottom part of Fig. 3. The correspondence between the protein stability calculations so far described and the neural progenitor types can be summarized as

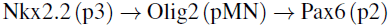

In the differentiation from pMN to p2, both Pax6 and Irx3 but not Olig2 are found in p2 (Dessaud et al., 2008; Balaskas et al., 2012). Moving away from the high-log *f*_O_2__ and log*a*_H_2_O_ limit of Fig. 3, Irx3 replaces Olig2 as the most stable proteins in the system. This pattern can be conceived as the product of irreversible apparent chemical reaction toward the most stable products in the system. Therefore, this reaction can be identified theoretically as the first step in an *unloading* stage, as portrayed in the upper part of Fig. 3. With further addition of Shh into the system, not only do its effects span a greater distance, but the system has more time to react, and to relax in response to the initial loading and formation of a non-equilibrium state.

With these theoretical considerations in mind, the calculated energies shown in Fig. 3 can be used to describe the rest of the major reactions. As noted elsewhere (Jessell, 2000; Dessaud et al., 2008), the transition from p2 to p1 is accompanied by the loss of Nkx6.1 and the appearance of Dbx2. In Fig. 3, decreasing log *f*_O_2__ and log*a*_H_2_O_ favors formation of Dbx2 relative to Nkx6.1. The appearance of Nkx6.2 after Irx3 is also characteristic of differentiation of p1 from p2 (Table 1); this reaction can be located as well in Fig. 3, at somewhat greater log *f*_O_2__ and log*a*_H_2_O_ values than the Nkx6.1 - Dbx2 reaction. Finally, at considerably lower values of log *f*_O_2__ and log*a*_H_2_O_, Fig. 3 shows the reaction Nkx6.2 - Dbx1, corresponding to the transition from p1 to p0 (Dessaud et al., 2008). In summary, the correspondence between the calculated relative stabilities of the proteins in the unloading stage of the model and the observed neural differentiation patterns can be represented as

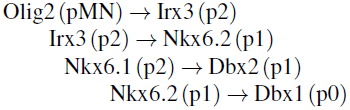

(Here, the staggered positions of the reactions represent the conditions at which the calculated stability transition occurs: lower log *f*_O_2__ to the right.)

The energetic analysis suggests a final, third stage, corresponding to conditions of maximal relative stability of Shh compared to the TFs (far right of Fig. 3). The overall greater stability of Shh compared to any TF at low log *f*_O_2__ and log*a*_H_2_O_ implies that Shh may reach metastable saturation in the tissue. Both the sequence of irreversible reactions leading to this point, and the loss of reaction potential at saturation, are thermodynamic constraints that may operate in conjunction with mechanistically based signaling processes in the developing embryo.

### Intracellular model: Interaction of Shh with cell-surface receptors and signaling of transcriptional activators

The expression of the transcription factors in different cells (Table 1) is brought about through a series of signals involving cell-surface receptors and intracellular transcriptional activators. The following paragraph is a very abridged description of the detailed signaling mechanism outlined in review papers (Ingham and McMahon, 2001; Wilson and Chuang, 2010; Hui and Angers, 2011), some aspects of which are explored further in the Discussion.

The intercellular morphogen Shh is a ligand that binds to the membrane receptor Patched (Ptch1). Upon binding, the Shh-Ptch1 complex is brought into the cell interior through endocytosis. The Shh-Ptch complex can be transported to recycling endosomes, from which Ptch1 is returned to the membrane, or transferred to lysosomes, where the proteins are degraded. When at the membrane, Ptch1 is a repressor of the transmembrane protein Smoothened (Smo). The removal of Ptch1 from the membrane leads to activation of Smo. Activated Smo transmits a signal to the DNA-binding Gli proteins (Gli1, Gli2, Gli3). Differential signaling events promote phosphorylation of Gli proteins, leading to their proteolysis and the formation of truncated sequences. The full-length forms are transcriptional activators (GliA), and the truncated forms are repressors (GliR). All of the Gli proteins can serve as activators, but the most important functions are activation by full-length Gli2 (Gli2A) and repression by truncated Gli3 (Gli3R). Ultimately, DNA binding by Gli modulates the expression of the downstream TFs, which in their turn bind DNA to activate the development-specific genes that determine the cellular identity.

The currently unknown mechanisms for repression of Smo by Ptch, activation of Gli by Smo, and control of downstream transcriptional events by GliR and GliA (Wilson and Chuang, 2010; Hui and Angers, 2011) constitute major gaps in the “relay model” for intracellular signal transduction summarized above. If we hypothesize that the downstream transcription factors (TFs; Table 1; Fig. 1) are formed through localized translation that is materially dependent on the lysosomal degradation of Shh and Ptch, then the expression of the TFs would be subject to thermodynamic constraints that can be theoretically assessed as described below. These intersection of these non-mechanistic constraints with known signaling mechanisms may help to resolve the open questions about the overall operation of the signaling process.

The compositional and energetic relations for the major interacting intracellular proteins are portrayed in the stability diagrams in Fig. 4. The methods are similar to the previous set of calculations, except that the reaction coefficients used in making Fig. 4 are normalized by protein length, so that the lines indicate equal activities of the proteins, not the residues (see Dick and Shock, 2011). The importance of normalization can be seen in the effects it has on the appearance of stability fields of proteins of different length. If the reactions in Figs. 1a-c were normalized (they were not), the size of the Shh field would grow; Shh is the shortest protein in that system. The most noticeable change there would be the shift of the Shh-Dbx1 boundary closer to log*a*_H_2_O_ = 0. In contrast, if the reactions in Figs. 4a-c (see below) were *not* normalized, the stability field for Gli3R, the shortest protein in that system, would not appear anywhere in the range of conditions shown on the diagram. Gli3R is derived from, and is much shorter than the activator form; it is thought that the relative abundances of Gli3R and Gli3A have a major control on the transcriptional response (Wilson and Chuang, 2010). Therefore, normalization is a desirable strategy as it enables theoretical prediction of the conditions where the proteins are equally abundant in metastable equilibrium (the field boundaries in Fig. 4b).

**Figure 4.**
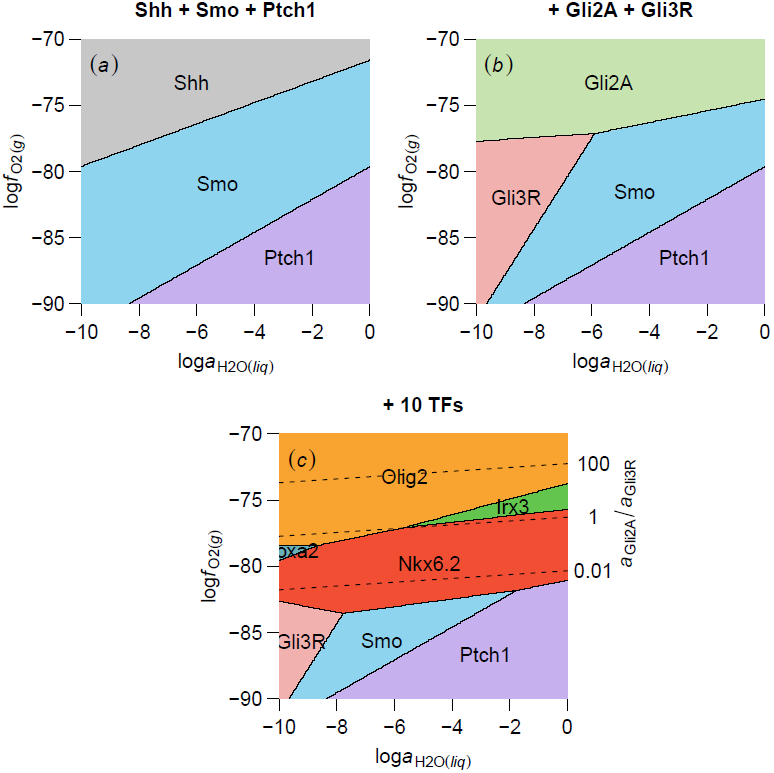
Theoretical metastability relations as a function of log*a*_H_2_O_ and log *f*_O_2__ for (*a*) Shh and its cell-surface receptor Patched 1 (Ptch1) and transmembrane protein Smoothened (Smo) and (*b*) the intracellular full-length transcriptional activator, Gli2A and truncated repressor, Gli3R and (*c*) downstream transcription factors controlled by Shh signaling. The dashed lines in (*c*) indicate the metastable equilibrium activity ratios of Gli2A to Gli3R.

In Fig. 4a the stability field for Smo appears between Shh and Ptch1; that is, a combination of the latter two proteins is unstable with respect to Smo. Also, Shh is stabilized at relatively oxidizing conditions. This is consistent with a comparatively lower average oxidation state of carbon in Smo and Ptch1, a common trend for membrane-associated proteins (Dick, 2014). In Fig. 4b, Gli2A (full-length activator) and Gli3R (truncated repressor) are added to the calculation. The sequence for Gli3R corresponds to residues G106 to E236 from the full-length Gli3 (Tsanev et al., 2009). The stability fields of Gli2A, and to a lesser extent Gli3R (at the lower end of the range of log*a*_H_2_O_ shown), replace the Shh field. In Fig. 4c, the downstream transcription factors are added (same as in Figs. 1a-c). The fields for Gli2A and, to some extent, Gli3R are covered by those of the transcription factors; showing that the latter are more stable than the former. The log *f*_O_2__-log*a*_H_2_O_ values corresponding to different ratios of chemical activities (concentrations) of Gli2A/Gli3R are overlaid in Fig. 4c to suggest possible downstream control by redox buffering (see below).

These chemical thermodynamic relationships have some conceptual implications that can be compared with observations in cell biology. *In the model*, the greater stability of Smo compared to Shh+Ptch1 in Fig. 4a is compatible with the notion that degradation of Shh+Ptch1 could lead to increased formation of Smo. *In the cell*, Shh binding to Ptch1 is followed by endocytosis of the Shh+Ptch1 complex and targeting to the lysosome for degradation (Ingham and McMahon, 2001). Shh binding also removes the inhibition by Ptch on Smo that promotes the turnover of Smo and prevents its accumulation at the membrane (Wilson and Chuang, 2010). *In the model*, Gli2A and Gli3R are together more stable than Shh. Therefore, addition of Shh to the system increases the potential for formation of Gli2A and Gli3R, and a greater proportion of Shh relative to more reduced proteins (such as the receptor Ptch) would lead to a greater potential for formation of Gli2A relative to Gli3R. *In the cell*, activation of Shh signaling leads to formation of Gli2A and a decrease in the half-life of Gli3 (Hui and Angers, 2011). *In the model*, the relatively low slope of the Gli2A-Gli3R reaction boundary reflects a low sensitivity to changing log*a*_H_2_O_ and correspondingly strong connection to changes in oxidation state. If it is supposed that Gli2A/Gli3R concentrations are sufficient to act as a redox buffer (Fig. 4c), it gives differences in the relative potential for formation of the different transcription factors, through which a selective force may be imparted on protein translation. *In the cell*, the actual mechanism of transmitting the signal from Gli to downstream gene products is unknown, but it is thought to have an essential dependence on the Gli concentrations and GliA/GliR ratios (Wilson and Chuang, 2010; Hui and Angers, 2011).

### Comparing the intercellular and intracellular models

Note the occurrence of stepwise patterns in the metastability relations of proteins in both the intracellular and intercellular models. Figs. 4a-c show, in order of increasing stability,

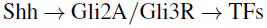

The overall cellular protein expression patterns during signal transduction (see above) corroborate this result. This ordering is not a contrived result based on an arbitrary ordering of system definitions but necessarily follows from the relative energies of the proteins, as was shown for the transcription factors in Figs. 1 and 2.

A progressive transition between energy levels is also apparent in the intercellular model. Consider the point in Fig. 3 where Olig2 becomes more stable than Nkx2.2. As a simplification of the loading-unloading model described above, if the system is allowed to progress toward metastable equilibrium under constant log *f*_O_2__ and log*a*_H_2_O_, the next most stable proteins would be Dbx1, Nkx6.1, Irx3, and Nkx6.2, in that order. Under additional assumptions, this stability ranking gives

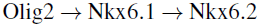

which closely follows the pMN to p2 to p1 progression (Table 1). The additional assumptions needed for this result, i.e. that Dbx1 and Irx3 are meta-metastable, are useful findings in themselves, as they may guide the formation of hypotheses about additional, extra-thermodynamic sources of specificity that could arise from lower-level signaling mechanisms.

In both of the immediately preceding examples, as the system is approaching metastable equilibrium, it does not immediately go to the lowest-energy state, but instead steps through the energy levels. This is reminiscent of the the Ostwald step rule, known mostly in the context of mineral precipitation (Morse and Casey, 1988). In some systems, it is not the most stable mineral that precipitates first, but a less stable one that is closer in energy to the nucleation event. The sequential progression of a system through close-to-equilibrium steps can result in lower entropy production than a reaction that immediately forms the most stable species (van Santen, 1984). Thus, the apparent stepwise energetic relations might constitute evidence for processes that operate in accordance with the minimum entropy production principles for near-equilibrium systems (Prigogine and Nicolis, 1971).

## DISCUSSION

Some of the most pressing questions in developmental biology focus on the difference between descriptive and explanatory accounts, and the levels of organization at which explanation is relevant (Minelli and Pradeu, 2014). The model presented above, not being mechanistic in nature, is conceived as a high-level descriptive account, with an emphasis on the recognition of overall patterns. Compared to conventional representations of signaling through molecular interactions, binding and gene activation (“relay models”), this approach uses different state variables and looks at a different “phase space” for the system. The model provides for the construction of specific, quantitative descriptions based on assumed organizational constraints; therefore the present model provides a framework (or perhaps “reference scheme”; Gould, 1980) for describing the system.

Despite the ongoing efforts to elucidate the mechanistic steps of signaling interactions, there are many unclear connections and significant questions (Wilson and Chuang, 2010; Hui and Angers, 2011). By what mechanisms does Ptch repress the activity of Smo? How are Shh concentrations passed through the intracellular signaling apparatus (“interpreted”)? How are “combinations of Ci/Gli activators and repressors within a given cell … utilized to produce a specific transcriptional response” (Wilson and Chuang, 2010)?

The signaling starting with Shh, through activation of Smo and Gli, is thought to result in transcriptional response that leads to the synthesis of the downstream transcription factors (Pax6, Olig2, etc.). However, there may be relevant control processes beyond the transcriptional stage. The processes of mRNA transport and localized translation have emerged as a nexus for response to external stimuli in mature neurons (Jung et al., 2014). Many different mechanisms of translation initiation have been identified, but as with the uncertainty about specificity of transcriptional signaling, many questions still remain about translational control (Jung et al., 2014).

In addition to the properties of network interactions, including feedback, the behavior of a system also depends on restraints, i.e. the environment (Noble, 2010). Accounting for the integration of signaling systems with the spatial organization of the cell, including subcellular microenvironments, may require recognition of forces acting at higher levels, which might not be revealed through dissection and analysis of mechanisms. The thermodynamic model provides such a high-level description: starting with biologically informed assumptions about the sets of locally interacting proteins, the calculations proceed from mass balance and energetic considerations to give predictions – in fact, statements of probability – about the formation and degradation of proteins. The model carries no specific mechanistic dependencies, but can be formulated in terms of chemical potentials that characterize possible local equilibrium states of subcellular microenvironments.

In many pictures of endocytic pathways that contribute to intracellular signaling, the passage of endocytosed material to the lysosome (i.e. heterolysosome, as it contains exogenous material; de Duve and Wattiaux, 1966) is shown as a dead end. Lysosomal degradation is often regarded as the termination of the signal, but the degradation products certainly do not simply vanish. The degradation products of lysosomes are amino acids and other small molecules that go back into production of biomacromolecules (Levine, 2007). There is currently little experimental evidence for a direct material link between the endocytic pathway and protein synthesis. However, Blower (2013) raised the possibility that “mRNAs linked to endosomes could couple activation of a cell surface receptor to a translational response”. Apart from endosomal activity, a correlation between location of ubiquitin-dependent proteolysis and translation has been noted (Pines and Lindon, 2005).

These bits of evidence from cellular studies combined with the requirements of mass balance can be used to suggest the hypothesis that the degradation products of endocytosed ligand-receptor complexes contribute to the localized synthesis of other signaling proteins (such as Gli) or the downstream transcription factors. A corollary of this hypothesis is that thermodynamic constraints will be evident in the reaction products – the newly translated proteins. The thermodynamic model presented here quantifies these constraints in an ideal case. Shown in Fig. 5 is a sequence of cellular states representing the hypothesized formation of a cellular subsystem, which is a localized, partially closed environment that is fed by endocytosed proteins and in turn becomes the source of material for newly locally translated proteins.

**Figure 5.**
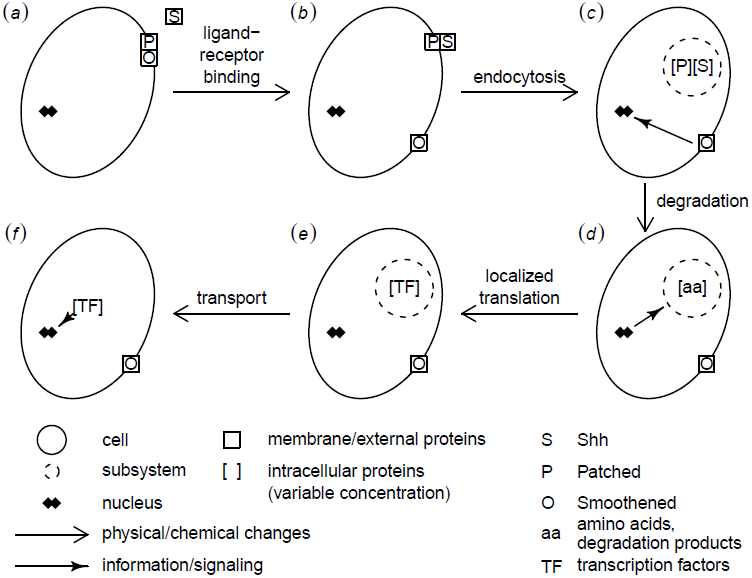
Conceptual model for the formation of transcription factors that are induced by Shh signaling. The filled arrows indicate intracellular information transfer associated with classical molecular mechanisms of signal transduction and gene expression. A partially closed cellular subsystem derived from endocytosed material (broken circles) is hypothesized to be the source of material for the localized translation of proteins. The corresponding material constraints are explored theoretically using the thermodynamic model described in this study.

The diagram also shows the general context of the molecular relay model of signal transduction leading to gene regulation (e.g. the arrow leading from Smo to the nucleus in Fig. 5c). Of course, the “phenomena at the macroscopic level must be reconciled with interactions on the level of molecular interactions” (Drack and Wolkenhauer, 2011). The overlapping possibility spaces of the relay model and the thermodynamic model would provide a more specific outcome than either type of process alone. This combination of perspectives can perhaps illuminate some of the open questions about the operation of the signaling network. For example, if the question about “specific transcriptional response” to varying GliA/GliR concentrations (Wilson and Chuang, 2010) is broadened to consider the possibility of localized degradation (of Gli) and translation, then the relative stabilities of the proteins suggests a role of macroscopic thermodynamic constraints in affecting the potential for formation of different transcription factors (Fig. 4c).

The physical balance of forces gives rise to constraints for operations within the whole organism, which Gould generalized as “structural integration” (Gould, 1980). These constraints of variation, as internal factors of evolution, are revealed in structure and development that follow the “laws of form” and the “transformation of the possible” (Gould, 1983, p. 144, p. 157). Thermodynamics is, after all, about limiting the possibilities. This leads to a new look at a fundamental question: Why do the proteins have the sequences they do? If we consider the chemical composition and energies of the proteins, the model described here showed plausible constraints on their limits of stability and order of formation. By analogy to Gould’s notion of structural integration, the constraints associated with chemical composition and energetics may be regarded as “chemical integration” within the organism. Organismal features constrained by structural integration are highly conserved in evolution: “Minor adaptations are multifarious, but major reorganizations are rare and strongly constrained by inherited structure” (Gould, 1980). The same may hold as well for developmental systems that depend on chemical integration of biomacromolecules. Thus, a high degree of chemical integration would tend to promote evolutionary conservation, and might contribute to the deep homology found between hedgehog morphogenetic systems in *Drosophila* and vertebrates (Gilbert et al., 1996).

The attributes of stability and equilibrium are closely associated with morphogenetic field concepts, of which morphogenetic gradients are a special case. Cellular determination starts with the condition of competence, which is a state of unstable equilibrium with multiple possible outcomes (Needham, 1942). The effect of an “organizer” is to provide a stimulus by which responding, competent cells irreversibly undergo determination. The specificity inherent in progression through differing degrees of stability is crucial; depending on the direction in which an unstable equilibrium is moved by the organizer, different outcomes will be selected, leading precisely to a differentiation (Rosen, 1968). A succession of organizational actions gives rise to the developmental or epigenetic landscape metaphor (Needham, 1942; Rosen, 1968). From these principles, a morphogenetic field can be defined that is a set of unstable entities (undifferentiated cells) in a spatial arrangement.

Needham’s “unstable equilibrium” (or, generally, an unstable steady state; see footnote 1) can be identified in our terms as a local equilibrium that is less stable than the metastable equilibrium – that is, any line below the locally highest line in Fig. 2. Take for example Shh: at highest log *f*_O2_ it is very unstable; as a chemical substance it has high potential to react. Depending on the direction and magnitude of change, the most stable protein will be Olig2, Irx3, Nkx6.2, Dbx2, or Shh - from highest to lowest log *f*_O_2__. The differential expression of these specific outcomes in terms of instability provides theoretical support for Shh to have “organizer” capabilities that, perhaps just as importantly, can be turned off when they are no longer needed (low log *f*_O_2__).

The study of redox dynamics may give further insight into signaling pathways. Whole-embryo redox potentials in zebrafish reported by Timme-Laragy et al. (2013) decrease steadily in the first ∼20 hours post fertilization, then fluctuate prior to hatching, rise during the hatching period (48-72 hpf), and achieve steady values in the early larva. A fall, then rise, in redox state is qualitatively consistent with the loading/unloading model depicted for the human proteins in Fig. 3. It would be interesting to compare the model with redox measurements in developmental systems at cellular or subcellular resolution, which are not currently available. Also, the model presented here is a simplification and many informative departures from this idealized state are possible. For example, a chemically integrated signaling system that is thermodynamically open may be regarded as being subject to cross talk with other redox-dependent metabolic pathways. In this situation, transformation of the protein molecules can be theoretically linked to compartment-specific subcellular redox states (Dick, 2014).

## CONCLUSION

Here I have outlined a high-level thermodynamic description of signaling and developmental patterns. There are reductionist assumptions inherent in either the conventional “relay models” or the descriptions based on chemical integration given here. However, by comparing patterns observed within different conceptual frameworks, complex biological properties might be distinguished from those that are simply emergent: “The whole is greater than the parts when these emergent properties cannot be explained solely by using properties that can be directly attributed to individuated parts” (Gilbert and Sarkar, 2000). A crucial distinction that may lead to a greater understanding of the whole is the explicit environmental dependence of macroscopic chemical thermodynamic models that, through Legendre transforms, utilize chemical potentials in the description of open systems. A comparable environmental dependence is operationally unavailable in current molecular-level explanations of signal transduction.

Chemical equilibrium is a theoretical point of departure from which we can construct quantitative descriptions of patterns in the physical world. In many areas of geochemistry, thermodynamic models are most useful not because they characterize stable equilibrium, but because by incorporating the concepts of affinity and metastability they provide the framework for identifying and quantifying non-equilibrium processes. This point was made by Anderson (2014): “Equilibrium thermodynamics cannot of course deal with irreversible reactions which disappear from thermodynamic state space … But thermodynamics can simulate such reactions using the affinity and the extent of reaction variable.” This has important practical benefits: “The study of metamorphic rocks shows that equilibrium has a broader meaning which is essential not only to the interpretation of rock history but also to understanding how thermodynamics deals with irreversible processes.”

Perhaps greater attention to this broader meaning of equilibrium will give more insight into biological processes as well, particularly those developmental processes that are characterized by irreversible steps of differentiation. Chemical thermodynamic models provide a high-level framework for describing compositional and energetic patterns related to differentiation, and foster novel hypotheses about linkages between subcellular microenvironments and macromolecular degradation and synthesis.

## Footnotes

1 A steady state is constancy in an open or dynamic system. Thermodynamic equilibrium is a condition of maximum entropy and minimum energy, that is, greater stability. Chemical thermodynamic equilibrium is a dynamic equilibrium in which the rates of forward and back reactions are equal (as a simple example for one reaction); therefore, chemical equilibrium is a special case of a steady state. Many authors use “equilibrium” when the more general notion of “steady state” is more appropriate; this and other ambiguities of terminology (Keller, 2008) apply to some of the citations in the following paragraphs. Any discussion or representation of chemical equilibrium, such as in this study, necessarily implies reference to a steady-state, so the following historical notes remain relevant. For example, the “dynamic equilibrium” of Schoenheimer is used in the general sense of steady state, without specifying thermodynamic equilibrium. Any manifestation of chemical equilibrium of proteins would be compatible with Schoenheimer’s dynamic equilibrium.

2 There are differing definitions of “system”; in theoretical terms it can be defined by the identities of the components and the bulk (elemental) composition, but in a more limited fashion as used here it is specified as well by the set of possible species, enforcing the metastable equilibrium assumptions. These theoretical systems are only approximate models for a real system.

3 The code used in this study depends on a recent version of CHNOSZ (version 1.0.3-13 at the time of writing), available at https://r-forge.r-project.org/projects/chnosz/

## REFERENCES

Alberty, R. A. (2003). Thermodynamics of Biochemical Reactions. John Wiley & Sons, Hoboken, New Jersey.

Anderson, G. M. (2005). Thermodynamics of Natural Systems. Cambridge University Press, 2nd edition.

Anderson, G. M. (2014). The entropy paradox and overstepping in metamorphic reactions. Chemical Geology, 384(0):10–15. DOI 10.1016/j.chemgeo.2014.06.020.

Balaskas, N., Ribeiro, A., Panovska, J., Dessaud, E., Sasai, N., Page, K., Briscoe, J., and Ribes, V. (2012). Gene regulatory logic for reading the Sonic Hedgehog signaling gradient in the vertebrate neural tube. Cell, 148(1):273–284. DOI 10.1016/j.cell.2011.10.047.

Bizzarri, M., Palombo, A., and Cucina, A. (2013). Theoretical aspects of Systems Biology. Progress in Biophysics and Molecular Biology, 112(1-2):33–43. DOI 10.1016/j.pbiomolbio.2013.03.019.

Blower, M. D. (2013). Molecular insights into intracellular RNA localization. In Jeon, K. W., editor, International Review of Cell and Molecular Biology, volume 302, pages 1–39. Academic Press. DOI 10.1016/B978-0-12-407699-0.00001-7.

Ciechanover, A. (2013). Intracellular protein degradation: from a vague idea through the lysosome and the ubiquitin-proteasome system and onto human diseases and drug targeting. Bioorganic & Medicinal Chemistry, 21(12):3400–3410. DOI 10.1016/j.bmc.2013.01.056.

Cohen, M., Page, K. M., Perez-Carrasco, R., Barnes, C. P., and Briscoe, J. (2014). A theoretical framework for the regulation of Shh morphogen-controlled gene expression. Development, 141(20):3868–3878. DOI 10.1242/dev.112573.

De Donder, Th. and Van Rysselberghe, P. (1936). Thermodynamic Theory of Affinity. Stanford University Press.

de Duve, C. and Wattiaux, R. (1966). Functions of lysosomes. Annual Review of Physiology, 28:435–492. DOI 10.1146/annurev.ph.28.030166.002251.

Dessaud, E., McMahon, A. P., and Briscoe, J. (2008). Pattern formation in the vertebrate neural tube: a sonic hedgehog morphogen-regulated transcriptional network. Development, 135(15):2489–2503. DOI 10.1242/dev.009324.

Dick, J. M. (2008). Calculation of the relative metastabilities of proteins using the CHNOSZ software package. Geochemical Transactions, 9:10. DOI 10.1186/1467-4866-9-10.

Dick, J. M. (2014). Average oxidation state of carbon in proteins. Journal of the Royal Society Interface, 11:20131095. DOI 10.1098/rsif.2013.1095.

Dick, J. M., LaRowe, D. E., and Helgeson, H. C. (2006). Temperature, pressure, and electrochemical constraints on protein speciation: group additivity calculation of the standard molal thermodynamic properties of ionized unfolded proteins. Biogeosciences, 3(3):311–336. DOI 10.5194/bg-3-311-2006.

Dick, J. M. and Shock, E. L. (2011). Calculation of the relative chemical stabilities of proteins as a function of temperature and redox chemistry in a hot spring. PLoS ONE, 6(8):e22782. DOI 10.1371/journal.pone.0022782.

Drack, M. and Wolkenhauer, O. (2011). System approaches of Weiss and Bertalanffy and their relevance for systems biology today. Seminars in Cancer Biology, 21(3):150–155. DOI 10.1016/j.semcancer.2011.05.001.

Garcia, H. G., Kondev, J., Orme, N., Theriot, J. A., and Phillips, R. (2011). Thermodynamics of biological processes. In Biothermodynamics, volume 492D of Methods In Enzymology, pages 27–59. Elsevier. DOI 10.1016/B978-0-12-381268-1.00014-8.

Gibbs, J. W. (1875). On the equilibrium of heterogeneous substances (first part). Transactions of the Connecticut Academy of Arts and Sciences, 3:108–248.

Gilbert, S. F. (1996). Enzymatic adaptation and the entrance of molecular biology into embryology. In Sarkar, S., editor, The Philosophy and History of Molecular Biology: New Perspectives, pages 101–123. Kluwer Academic Publishers, Dordrecht.

Gilbert, S. F., Opitz, J. M., and Raff, R. A. (1996). Resynthesizing evolutionary and developmental biology. Developmental Biology, 173(2):357–372. DOI 10.1006/dbio.1996.0032.

Gilbert, S. F. and Sarkar, S. (2000). Embracing complexity: organicism for the 21st century. Developmental Dynamics, 219(1):1–9. DOI 10.1002/1097-0177(2000)9999:9999<::AID-DVDY1036>3.0.CO;2-A.

Gould, S. J. (1971). D’Arcy Thompson and the science of form. New Literary History, 2(2):229–258.

Gould, S. J. (1980). The evolutionary biology of constraint. Daedalus, 109(2):39–52.

Gould, S. J. (1983). Hen’s Teeth and Horse’s Toes. W. W. Norton & Company, New York.

Green, S. and Wolkenhauer, O. (2013). Tracing organizing principles: learning from the history of systems biology. History and Philosophy of the Life Sciences, 35:553–576.

Helgeson, H. C. (1967). Solution chemistry and metamorphism. In Abelson, P. H., editor, Researches in Geochemistry, volume 2, pages 362–404. Wiley, New York.

Helgeson, H. C. (1970). Description and interpretation of phase relations in geochemical processes involving aqueous solutions. American Journal of Science, 268(5):415–438. DOI 10.2475/ajs.268.5.415.

Helgeson, H. C. (1979). Mass transfer among minerals and hydrothermal solutions. In Barnes, H. L., editor, Geochemistry of Hydrothermal Ore Deposits, pages 568–610. Wiley, New York, 2nd edition.

Helgeson, H. C., Garrels, R. M., and Mackenzie, F. T. (1969). Evaluation of irreversible reactions in geochemical processes involving minerals and aqueous solutions. II. Applications. Geochimica et Cosmochimica Acta, 33(4):455–481. DOI 10.1016/0016-7037(69)90127-6.

Helgeson, H. C., Knox, A. M., Owens, C. E., and Shock, E. L. (1993). Petroleum, oil-field waters, and authigenic mineral assemblages: are they in metastable equilibrium in hydrocarbon reservoirs? Geochimica et Cosmochimica Acta, 57(14):3295–3339. DOI 10.1016/0016-7037(93)90541-4.

Hui, C.-c. and Angers, S. (2011). Gli proteins in development and disease. Annual Review of Cell and Developmental Biology, 27(1):513–537. DOI 10.1146/annurev-cellbio-092910-154048. PMID: 21801010.

Ingham, P. W. and McMahon, A. P. (2001). Hedgehog signaling in animal development: paradigms and principles. Genes and Development, 15(23):3059–3087. DOI 10.1101/gad.938601.

Jessell, T. M. (2000). Neuronal specification in the spinal cord: inductive signals and transcriptional codes. Nature Reviews. Genetics, 1(1):20–29. DOI 10.1038/35049541.

Judson, H. F. (1979). The Eighth Day of Creation. Cold Spring Harbor Laboratory Press, New York. (Commemorative Edition, 2013).

Jung, H., Gkogkas, C. G., Sonenberg, N., and Holt, C. E. (2014). Remote control of gene function by local translation. Cell, 157(1):26–40. DOI 10.1016/j.cell.2014.03.005.

Keller, E. F. (2008). Organisms, machines, and thunderstorms: A history of self-organization, part one. Historical Studies In the Natural Sciences, 38(1):45–75. DOI 10.1525/hsns.2008.38.I.45.

Kitcher, P. (1999). The hegemony of molecular biology. Biology and Philosophy, 14(2):195–210. DOI 10.1023/A:1006686417669.

Korzhinskii, D. S. (1965). The theory of systems with perfectly mobile components and processes of mineral formation. American Journal of Science, 263(3):193–205. DOI 10.2475/ajs.263.3.193.

Lander, A. D. (2007). Morpheus unbound: reimagining the morphogen gradient. Cell, 128(2):245–256. DOI 10.1016/j.cell.2007.01.004.

LaRowe, D. E. and Dick, J. M. (2012). Calculation of the standard molal thermodynamic properties of crystalline peptides. Geochimica et Cosmochimica Acta, 80:70–91. DOI 10.1016/j.gca.2011.11.041.

Levine, B. (2007). Cell biology: autophagy and cancer. Nature, 446(7137):745–747. DOI 10.1038/446745a.

Minelli, A. and Pradeu, T. (2014). Theories of development in biology–problems and perspectives. In Minelli, A. and Pradeu, T., editors, Towards a Theory of Development, pages 1–14. Oxford University Press.

Morse, J. W. and Casey, W. H. (1988). Ostwald processes and mineral paragenesis in sediments. American Journal of Science, 288(6):537–560. DOI 10.2475/ajs.288.6.537.

Needham, J. (1942). Biochemistry and Morphogenesis. Cambridge University Press.

Noble, D. (2010). Biophysics and systems biology. Philosophical Transactions of the Royal Society A: Mathematical, Physical and Engineering Sciences, 368(1914):1125–1139. DOI 10.1098/rsta.2009.0245.

Pickett, S. T. A., Kolasa, J., and Jones, C. G. (2007). Ecological Understanding: The Nature of Theory and the Theory of Nature. Academic Press, 2nd edition.

Pines, J. and Lindon, C. (2005). Proteolysis: anytime, any place, anywhere? Nature Cell Biology, 7(8):731–735. DOI 10.1038/ncb0805-731.

Prigogine, I. and Nicolis, G. (1971). Biological order, structure and instabilities. Quarterly Reviews of Biophysics, 4(2-3):107–148. DOI 10.1017/S0033583500000615.

Rosen, R. (1968). Recent developments in the theory of control and regulation of cellular processes. In Bourne, G. H., Danielli, J. F., and Jeon, K. W., editors, International Review of Cytology, volume 23, pages 25–88. Academic Press. DOI 10.1016/S0074-7696(08)60269-7.

Scheiner, S. M. (2010). Toward a conceptual framework for biology. Quarterly Review of Biology, 85(3):293–318. DOI 10.1086/655117.

The UniProt Consortium (2015). UniProt: a hub for protein information. Nucleic Acids Research, 43(D1):D204–D212. DOI 10.1093/nar/gku989.

Thompson, Jr., J. B. (1959). Local equilibrium in metasomatic processes. In Abelson, P. H., editor, Researches in Geochemistry, pages 427–457. John Wiley & Sons, New York.

Timme-Laragy, A. R., Goldstone, J. V., Imhoff, B. R., Stegeman, J. J., Hahn, M. E., and Hansen, J. M. (2013). Glutathione redox dynamics and expression of glutathione-related genes in the developing embryo. Free Radical Biology and Medicine, 65(0):89–101. DOI 10.1016/j.freeradbiomed.2013.06.011.

Tsanev, R., Tiigimägi, P., Michelson, P., Metsis, M., Østerlund, T., and Kogerman, P. (2009). Identification of the gene transcription repressor domain of Gli3. FEBS Letters, 583(1):224–228. DOI 10.1016/j.febslet.2008.12.010.

van Santen, R. A. (1984). The Ostwald step rule. Journal of Physical Chemistry, 88(24):5768–5769. DOI 10.1021/j150668a002.

Waddington, C. H. (1956). Principles of Embryology. MacMillan, New York.

Wilson, C. W. and Chuang, P.-T. (2010). Mechanism and evolution of cytosolic Hedgehog signal transduction. Development, 137(13):2079–2094. DOI 10.1242/dev.045021.

